# Effects of potassium fertilizer on dry matter accumulation and potassium absorption characteristics of maize (Zea mays L.) inbred lines with different yield types

**DOI:** 10.1101/2021.04.06.438571

**Authors:** Ya-fang Fan, Ju-lin Gao, Ji-ying Sun, Jian Liu, Zhi-jun Su, Shu-ping Hu, Zhi-gang Wang, Xiao-fang Yu

## Abstract

In this experiment, 20 different maize inbred lines were used as basic materials, which were divided into high-yield type, middle-yield type and low-yield type by yield cluster analysis. A 3-year long-term positioning field experiment (2016–2018) was carried out to assess the effects of potassium fertilizer on dry matter, potassium absorption and potassium absorption efficiency of maize inbred lines with different yield types. The results showed that the dry matter, potassium absorption and potassium absorption efficiency of leaf, stem, bract, rachis and grain of maize inbred lines have been significantly increased with potassium application in comparison with no potassium application. Potassium fertilizer could improve the dry matter accumulation capacity and potassium absorption characteristics of maize, promote the potassium absorption by maize plants, influence the growth and development of maize, and then improve the maize yield. The decrease of dry matter, potassium absorption and potassium absorption efficiency of high-yield maize inbred lines was smaller under low potassium stress. The high-yield maize inbred lines had stronger ability to maintain higher dry matter accumulation and potassium absorption characteristics under low potassium stress. The study provided a theoretical basis for the breeding of potassium-efficient maize materials.

## Introduction

Maize is the largest food crop in China, which contributes more than 80% to the increase of grain production. It is the main force of cereal production and the key to ensure food security. The high quality and high yield of maize is of great significance to food security and stability [1, 2].

Potassium is an essential nutrient for the growth and development of maize, which is involved in many important physiological processes. Sufficient potassium helps to ensure high yield of maize [3–6]. Dry matter is the highest form of photosynthetic maize products, the accumulation of which is closely linked to the maize yield. A large number of field experiments showed that the increase in dry matter directly or indirectly promoted the increase in biomass and grain weight, which laid the foundation for the increase of yield [7, 8]. Dry matter was positively correlated with maize yield and components [9]. The soil fertility significantly affected dry matter of maize. Potassium application could promote the accumulation of dry matter and increase the yield of maize [10, 11]. The increase of yield was not only related to the accumulation of dry matter, but also to the absorption of potassium. Studies have shown that maize was necessary to absorb sufficient potassium to achieve a higher yield [12]. There has been a significant positive correlation between potassium absorption capacity and maize grain yield [13, 14], and the adequate potassium nutrition promoted the accumulation and absorption of potassium in the plant and then improved the yield of maize. Tan Deshui et al. [15] found that potassium fertilizer could significantly improve the maize yield through 13-year of continuous potassium application experiment.

In conclusion, the previous studies focused on the effects of potassium fertilizer on dry matter and potassium absorption characteristics of maize, but most of them concentrated on a specific variety or the same type of material, and few studies focused on different yield types of maize inbred lines. Therefore, this experiment was based on the premise of previous studies on the long-term positioning experiment of potassium fertilizer treatment with different yield types of maize inbred lines with genetic differences as experimental materials, and no potassium application was used as control. The differences in dry matter, potassium absorption and potassium absorption efficiency of different yield maize inbred lines were compared, and the response of different yield types of maize inbred lines to potassium fertilizer has been analyzed to select suitable maize inbred lines for rational utilization of potassium fertilizer resources, and to create a scientific basis for optimal management of potassium fertilizer resources.

## Materials and methods

### Site description

Field experiments were conducted from 2016 to 2018 at the Experimental Station of Inner Mongolia Agricultural University (40°33’N, 110°31’E), which is located in the middle and western of Inner Mongolia Autonomous Region, China. The 3-year average annual temperature was 22.1°C, the average annual sunshine hours were 1617.5h, and the average annual precipitation was 302.7ml, approximately 70% of the average annual precipitation occurred from June to September. The soil texture at the site was sandy loam (ISSS Classification, International Soil Science Society), with 19.35, 19.28 and 20.06 g kg^-1^ organic material, 61.36, 61.52 and 62.17 mg kg^-1^ available N, 5.46, 5.27 and 5.62 mg kg^-1^ available P, 93.65, 95.17 and 91.52 mg kg^-1^ available K, and the pH of 7.5,7.6 and 7.6, respectively in 2016, 2017 and 2018.

The field study was carried out on the official land which belonged to the key laboratory of crop cultivation and genetic improvement of Inner Mongolia Autonomous Region, permission was given after research application passing verification. During the field study none of endangered or protected species were involved. No specific permissions were required for conducting the field study because it was not carried out in protected area.

### Experimental materials

20 different maize inbred lines were selected as the test materials: FAPW, PHG39, LH51, K12, Shen 5003, NP49, HuangC, Jun92-8, Zi330, P2, LP5, 7810, IRF233, IB014, Ji63, G303, IRF236, M3401, L269, DH382.

### Experimental design and treatments

The experiment was carried out according to a split-plot design: potassium fertilizer levels were considered the main factors and the maize inbred lines were sub-plot factors, with three randomized replicates. The experiment set two potassium fertilizer levels, which K_2_O were 45 kg/hm^2^ and 0 kg/hm^2^, expressed as 45K and 0K. The planting density was 67500 plants ha^-1^. Each plot was 5 m long by 0.6 m wide and contained 8 rows, and each plot area measured 24m^2^. There were 40 plots in the experiment. The irrigation was carried out four times throughout the growing period and each quantity of irrigation was 750 m^3^ hm^-2^.Other management measures such as spraying and weeds were the same as field cultivation. Maize was sown in late April and harvested in early October during 2016-2018.

### Determination index and method

In the silking stage and maturity stage, maize plants were taken from each plot with three randomized replicates. In the silking stage, the maize plant was divided into two parts: leaf and stem, bract and rachis. At maturity, the maize plant was divided into three parts: leaf and stem, bract and rachis, grain. The plants were heated initially at 105°C for 30 min and oven-dried at 80°C to a constant weight before weighing. The dry matter was crushed and screened by 0.5 mm sieve and then decocted by using the H_2_SO_4_–H_2_O_2_ method [16]. The potassium content of maize was determined by flame photometers and the potassium absorption and efficiency were calculated.

At maturity, two rows (6m^2^) of maize in the middle of the test area were manually harvested of each plot to determine the grain yield, and weighed the grain weight after threshing. The moisture content of maize grain was measured by Grain Moisture Tester (PM-8188-a, Japan), and the grain yield was calculated [17].

### Date analysis

Microsoft Excel 2010 was used to input and arrange the data. IBM SPSS statistics 22.0 was used to finish the cluster analysis, ANOVA and correlation analysis on different maize inbred lines. LSD method was used to analyze the difference between treatments (P=0.05) [18]. Histograms were conducted using Sigma Plot 12.5. And Different letters on histograms indicated that means were statistically different at the 0.05 level. The data used in the charts and tables were the mean values of the data in 2016, 2017 and 2018.

## Results

### Cluster analysis of maize inbred lines under different potassium fertilizer levels

The cluster analysis of the maize yield was carried out at the level of 0K and 45K (Fig 1, Fig 2), and if the threshold value was set to 5, 20 maize inbred lines could be divided into three types under two potassium fertilizer levels. The first type was high-yield maize inbred line, including FAPW, PHG39, LH51, K12, Shen 5003 and NP49, which had higher yield under two different potassium levels. The range of yield was 6306.46~6600.69 kg/hm^2^ at 45K and 5820.87~6089.30 kg/hm^2^ at 0K. The second type was middle-yield maize inbred line, including Huang C, Jun 92-8, Zi 330, P2, LP5, 7810 and IRF233, which yield was in the middle under two different potassium levels. The range of yield was 4646.74~4892.28 kg/hm^2^ at 45K and 4412.02~4650.36 kg/hm^2^ at 0K. The third type was low-yield maize inbred line, including IB014, Ji63, G303, IRF236, M3401, L269 and DH382, which yield was lower under two different potassium levels. The range of yield was 3370.52~3567.51 kg/hm^2^ at 45K and 3280.46~3469.88 kg/hm^2^ at 0K.

**Fig 1.**
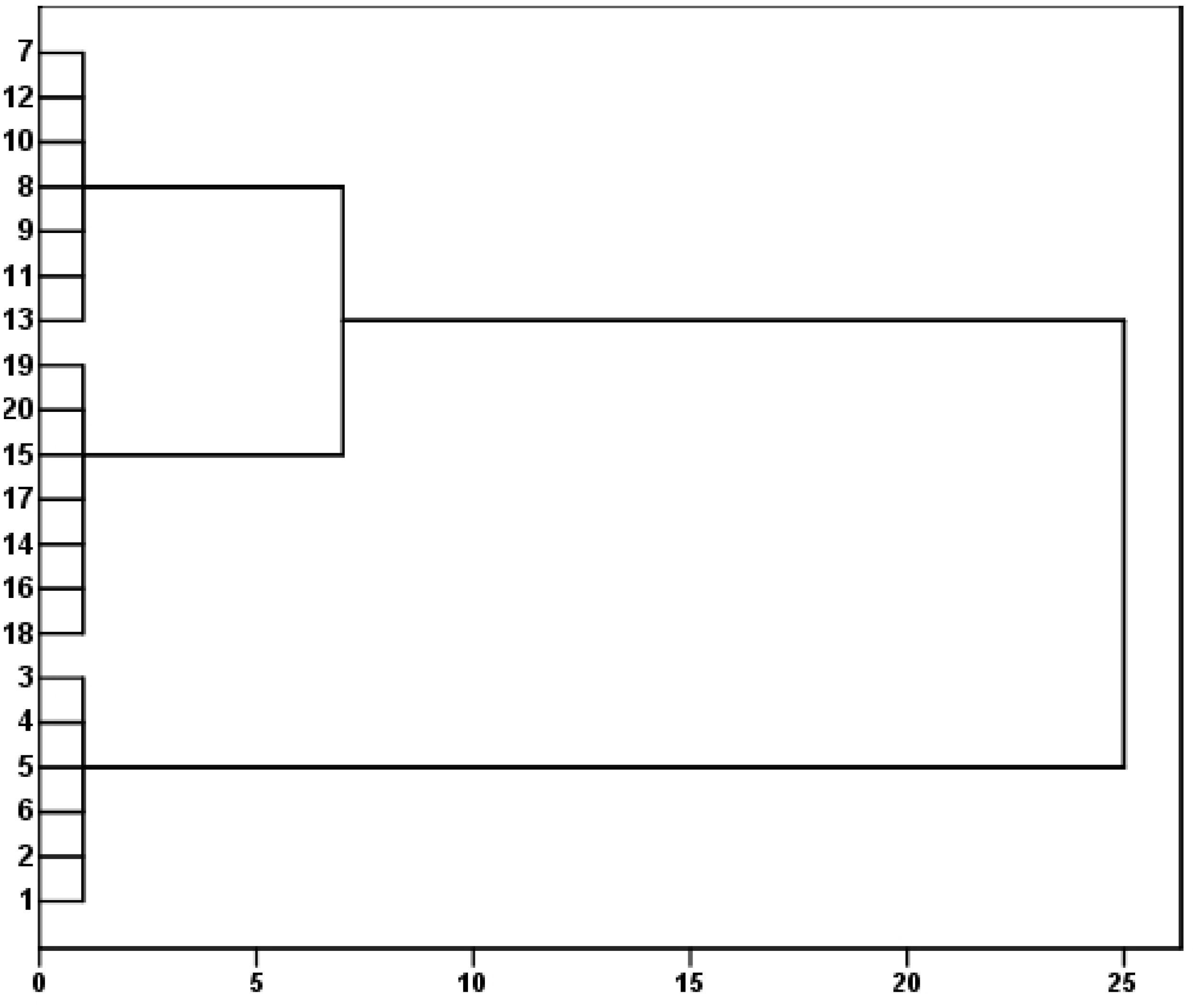
Cluster analysis of yield of maize inbred lines at 45K level Note: The abscissa represents the relative distance, and the ordinate represents 20 maize inbred lines, 1: FAPW; 2: PHG39; 3: LH51; 4: K12; 5: Shen5003; 6: NP49; 7: Huang C; 8: Jun92-8; 9: Zi330; 10: P2; 11: LP5; 12: 7810; 13: IRF233; 14: IB014; 15: Ji63; 16: G303; 17: IRF236; 18: M3401; 19: L269; 20: DH382. The same as below.

**Fig 2.**
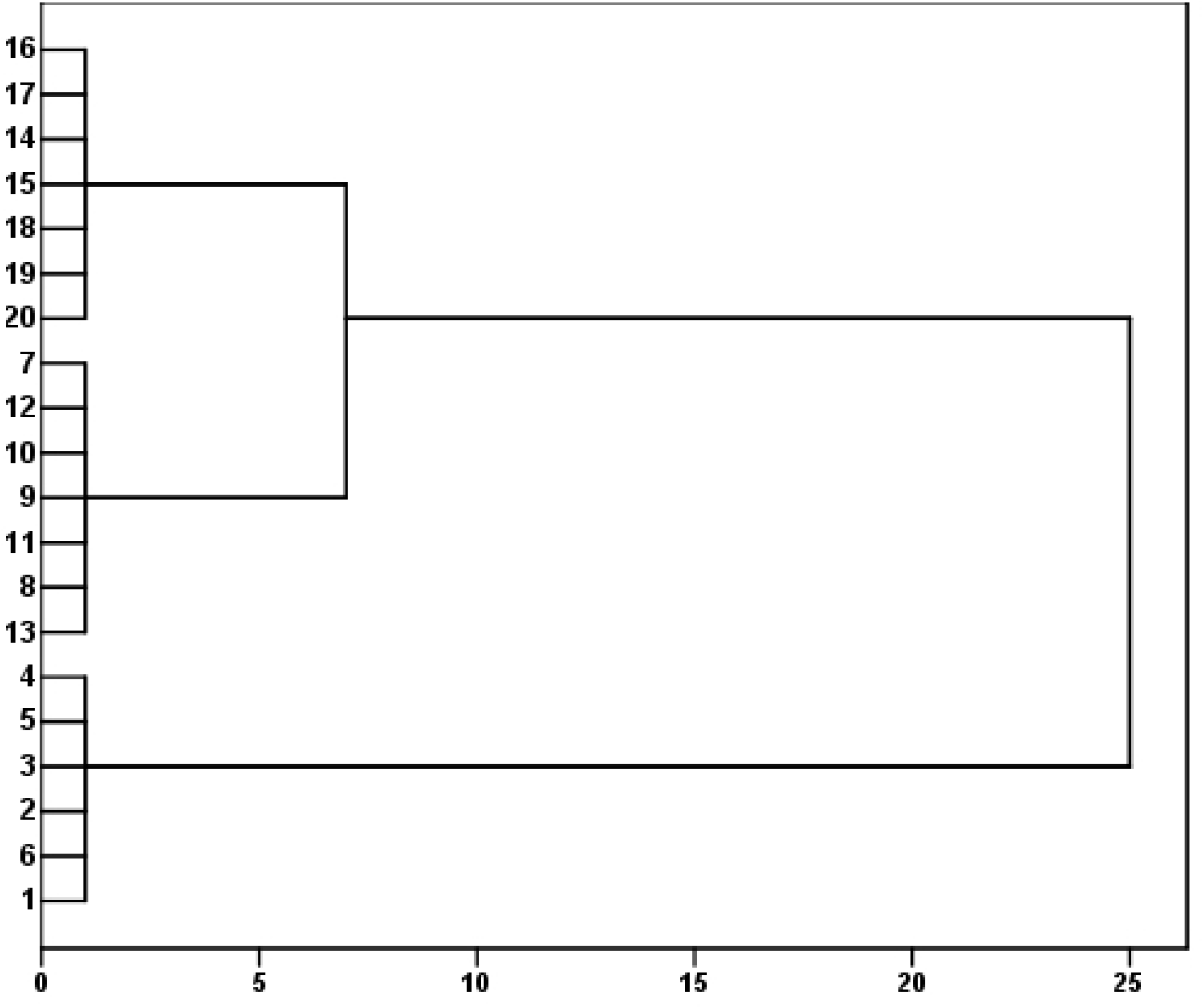
Cluster analysis of yield of maize inbred lines at 0K level

### Effects of potassium fertilizer on yield of maize inbred lines with different yield types

The grain weight, grain water content and yield of maize inbred lines varied significantly (p<0.05) between potassium levels and inbred lines, and the effects of interaction between inbred lines and potassium levels varied significantly (Table 1).

**Table 1.** ANOVA results for yield and related parameters of maize inbred lines

| Source of variation | DF | Grain weight per spike (g/fringe) | Water content (%) | Yield (kg/hm<sup>2</sup>) |
| --- | --- | --- | --- | --- |
| Potash application level (K) | 1 | 20.06** | 12.15** | 336027.80** |
| Yield type (T) | 2 | 3977.17** | 6.91** | 11422280.00** |
| K×T | 2 | 23.39** | 0.12** | 61447.11** |
| Error | 10 | 1.30 | 0.01 | 2593.35 |

Note: \*\* Significant at the 0.05 probability level. These values in the table were the mean squares (MS) of the ANOVA.

The yield of high-yield, middle-yield and low-yield type maize inbred lines increased by 7.96%~8.45%, 4.72%~5.35% and 2.41%~2.95% at 45K compared with that at 0K (Table 2). The yield of high-yield type maize inbred lines was 1.35 and 1.85 times higher than that of middle-yield and low-yield type at 45K, and 1.31 and 1.75 times higher than that of middle-yield and low-yield type at 0K. The yield of maize increased significantly under 45K, the yield increasing range of maize inbred lines with different yield types was as follows: high-yield type> middle-yield type>low-yield type.

**Table 2.**
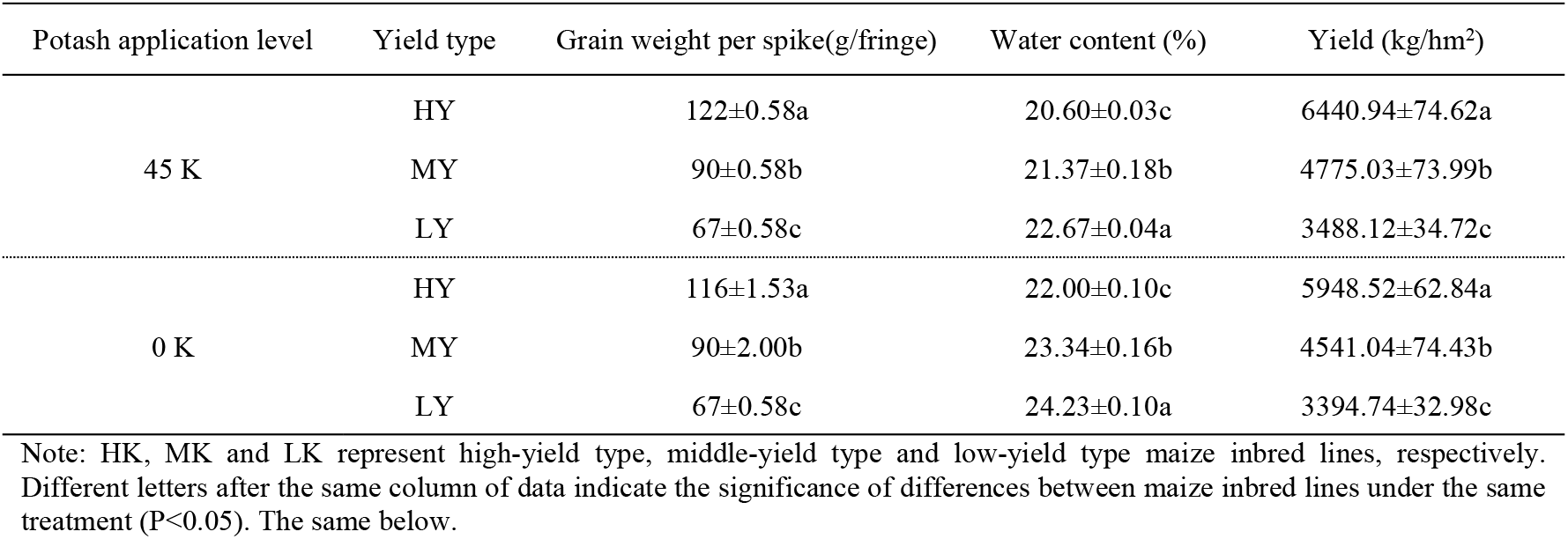
Effects of potassium fertilizer on yield and related parameters of maize inbred lines with different yield types

| Potash application level | Yield type | Grain weight per spike(g/fringe) | Water content (%) | Yield (kg/hm<sup>2</sup>) |
| --- | --- | --- | --- | --- |
| 45 K | HY | 122±0.58a | 20.60±0.03c | 6440.94±74.62a |
|  | MY | 90±0.58b | 21.37±0.18b | 4775.03±73.99b |
|  | LY | 67±0.58c | 22.67±0.04a | 3488.12±34.72c |
| 0 K | HY | 116±1.53a | 22.00±0.10c | 5948.52±62.84a |
|  | MY | 90±2.00b | 23.34±0.16b | 4541.04±74.43b |
|  | LY | 67±0.58c | 24.23±0.10a | 3394.74±32.98c |

Note: HK, MK and LK represent high-yield type, middle-yield type and low-yield type maize inbred lines, respectively. Different letters after the same column of data indicate the significance of differences between maize inbred lines under the same treatment ( $P<0.05$ ). The same below.

### Effects of potassium fertilizer on dry matter of maize inbred lines with different yield types

The dry matter of leaf and stem, bract and rachis, grain of maize inbred lines in the silking stage and maturity stage varied significantly (p<0.05) between potassium levels and inbred lines, and the effects of interaction between inbred lines and potassium levels varied significantly (Table 3).

**Table 3.** ANOVA results for dry matter of maize inbred lines

| Source of variation | DF | Dry matter in R1 |  | Dry matter in R6 |  |  |
| --- | --- | --- | --- | --- | --- | --- |
|  |  | Leaf and stem | Bract and rachis | Leaf and stem | Bract and rachis | Grain |
|  |  | (g/plant) | (g/plant) | (g/plant) | (g/plant) | (g/plant) |
| Potash application level (K) | 1 | 623.16** | 64.15** | 255.98** | 107.85** | 440.95** |
| Yield types (T) | 2 | 1472.05** | 446.37** | 1230.16** | 242.51** | 1826.14** |
| K×T | 2 | 31.13** | 5.93** | 24.35** | 9.62** | 42.20** |
| Error | 10 | 0.15 | 0.10 | 0.14 | 0.06 | 0.08 |

Note: R1 and R6 indicate the silking stage and maturity stage of maize respectively. The same below.

In the silking stage, the dry matter of leaf and stem of high-yield, middle-yield and low-yield type maize inbred lines increased by 18.41%~20.02%, 14.26%~16.31% and 11.58%~13.68% at 45K compared with that at 0K; the dry matter of bract and rachis increased by 14.41%~16.25%, 11.37%~13.47% and 6.95%~9.24%. The dry matter of leaf and stem of high-yield type maize inbred lines was 1.21 and 1.53 times higher than that of middle-yield and low-yield type at 45K, 1.18 and 1.44 times higher than that of middle-yield and low-yield type at 0K. The dry matter of bract and rachis of high-yield type maize inbred lines was 1.35 and 1.77 times higher than that of middle-yield and low-yield type at 45K, 1.31 and 1.66 times higher than that of middle-yield and low-yield type at 0K (Fig 3).

**Fig 3.**
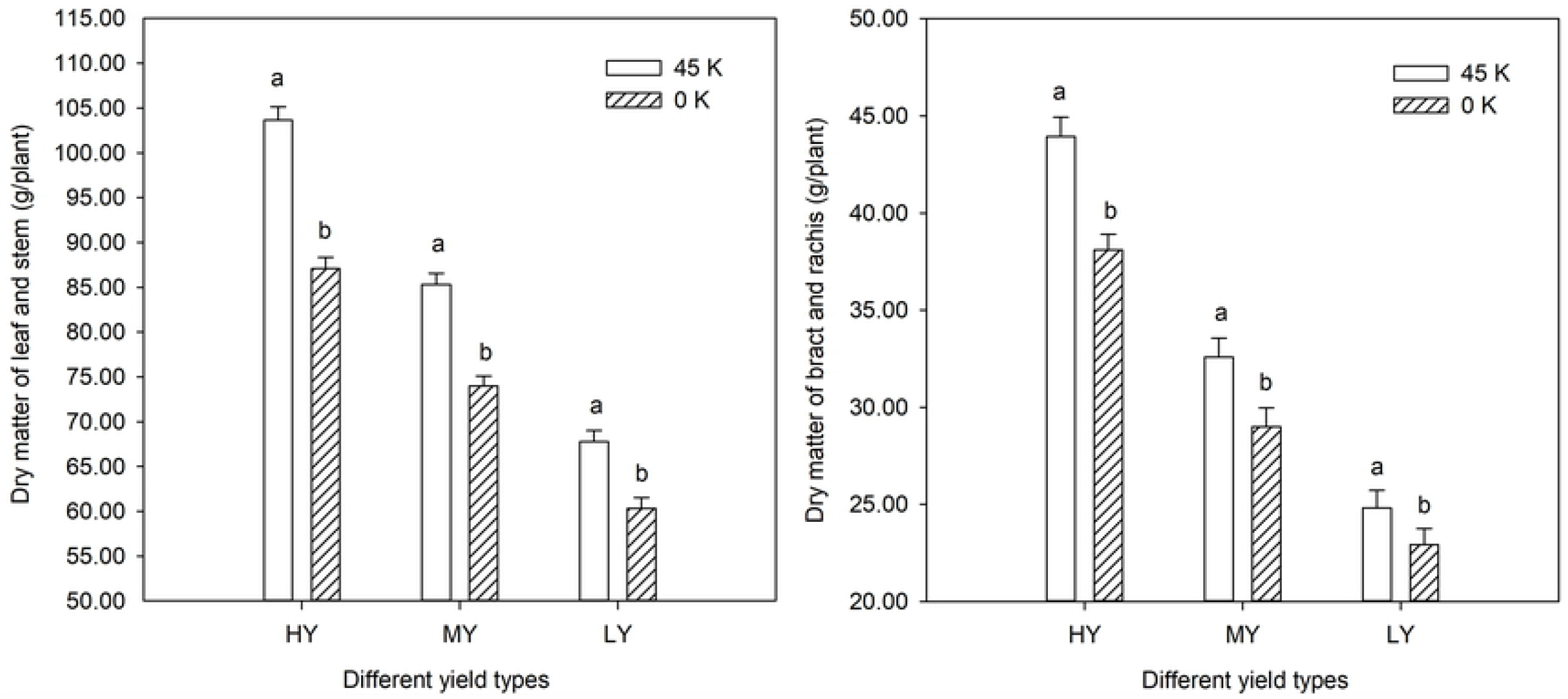
Effects of potassium fertilizer on dry matter of maize inbred lines with different yield types in the silking stage

At maturity, the dry matter of leaf and stem of high-yield, middle-yield and low-yield type maize inbred lines increased by 13.16%~14.84%, 9.16%~11.59% and 5.28%~6.96% at 45K compared with that at 0K; the dry matter of bract and rachis increased by 16.79%~18.99%, 11.38%~13.64% and 7.17%~9.70%; the dry matter of grain increased by 14.29%~15.23%, 9.58%~11.71% and 5.81%~7.63%. The dry matter of leaf and stem of high-yield type maize inbred lines was 1.29 and 1.50 times higher than that of middle-yield and low-yield type at 45K, 1.25 and 1.40 times higher than that of middle-yield and low-yield type at 0K. The dry matter of bract and rachis of high-yield type maize inbred lines was 1.21 and 1.43 times higher than that of middle-yield and low-yield type at 45K, 1.16 and 1.31 times higher than that of middle-yield and low-yield type at 0K. The dry matter of grain of high-yield type maize inbred lines was 1.21 and 1.51 times higher than that of middle-yield and low-yield type at 45K, 1.16 and 1.40 times higher than that of middle-yield and low-yield type at 0K (Fig 4).

The dry matter of maize inbred lines with different yield types were shown as follows: high-yield type>middle-yield type>low-yield type, and potassium fertilizer could significantly increase the dry matter of maize.

**Fig 4.**
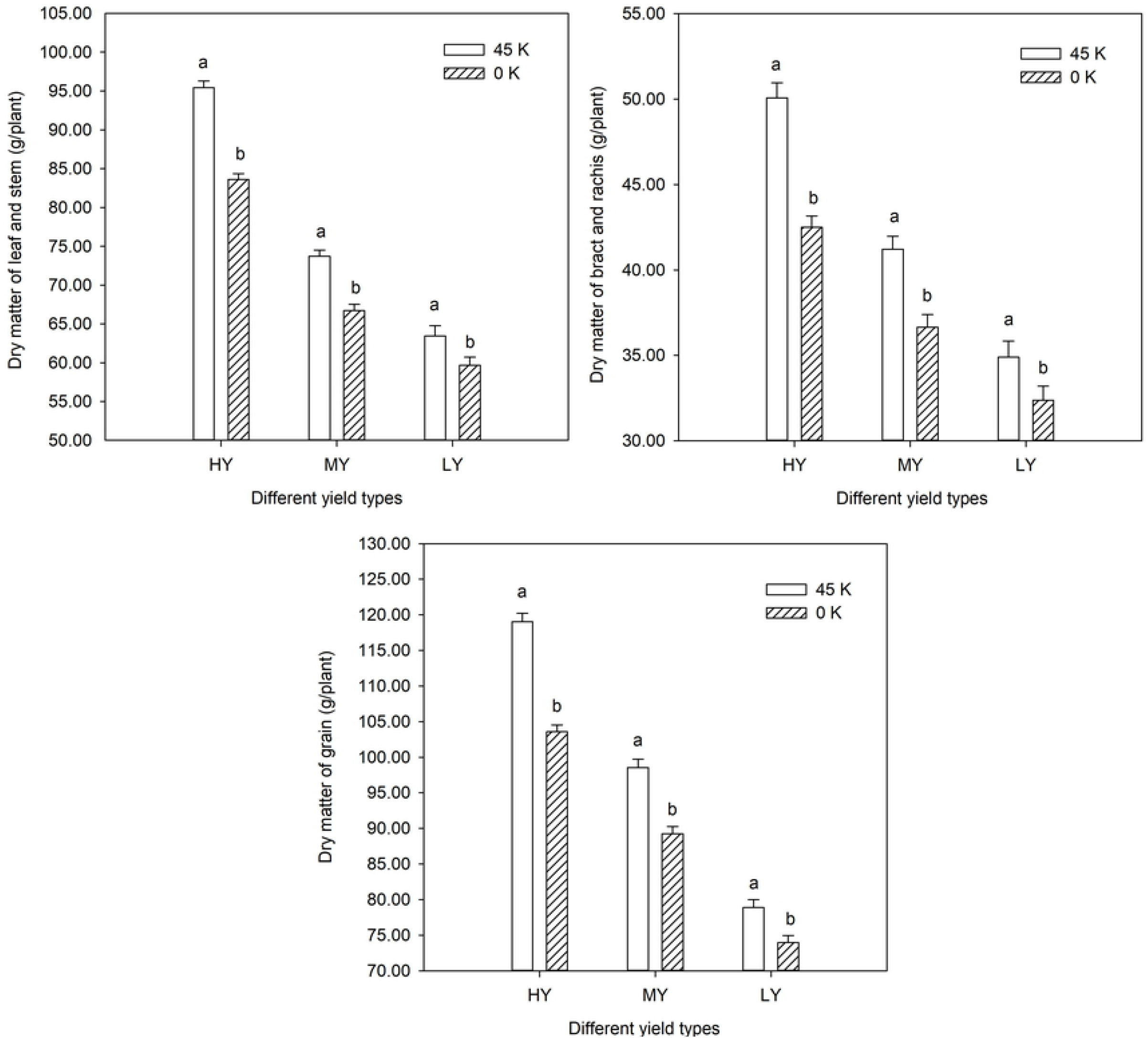
Effects of potassium fertilizer on dry matter of maize inbred lines with different yield types at maturity stage

### Effects of potassium fertilizer on potassium absorption of maize inbred lines with different yield types

The potassium absorption of leaf and stem, bract and rachis, grain of maize inbred lines in the silking stage and maturity stage varied significantly (p<0.05) between potassium levels and inbred lines, and the effects of interaction between inbred lines and potassium levels varied significantly (Table 4).

**Table 4.**
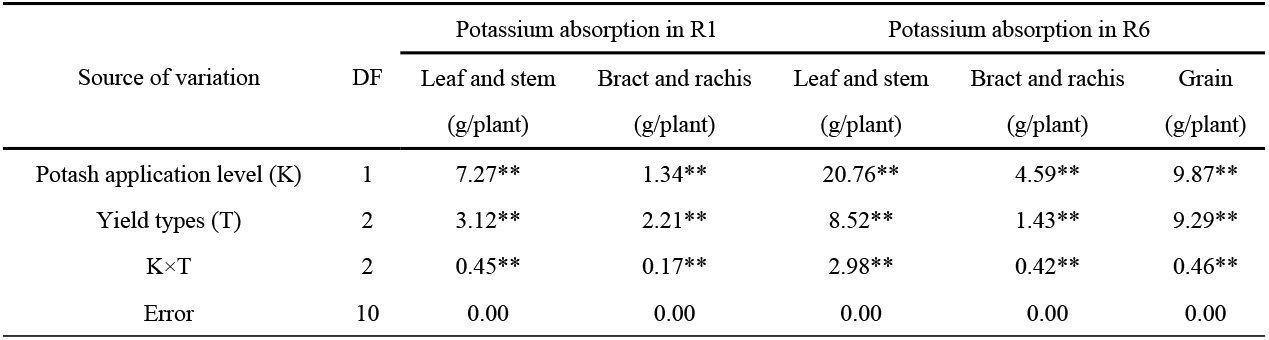
ANOVA results for potassium absorption of maize inbred lines

| Source of variation | DF | Potassium absorption in R1 |  | Potassium absorption in R6 |  |  |
| --- | --- | --- | --- | --- | --- | --- |
|  |  | Leaf and stem<br>(g/plant) | Bract and rachis<br>(g/plant) | Leaf and stem<br>(g/plant) | Bract and rachis<br>(g/plant) | Grain<br>(g/plant) |
| Potash application level (K) | 1 | 7.27** | 1.34** | 20.76** | 4.59** | 9.87** |
| Yield types (T) | 2 | 3.12** | 2.21** | 8.52** | 1.43** | 9.29** |
| K×T | 2 | 0.45** | 0.17** | 2.98** | 0.42** | 0.46** |
| Error | 10 | 0.00 | 0.00 | 0.00 | 0.00 | 0.00 |

In the silking stage, the potassium absorption of leaf and stem of high-yield, middle-yield and low-yield type maize inbred lines increased by 1.49~2.32 g/plant, 1.08~1.44 g/plant and 0.52~1.07 g/plant at 45K compared with that at 0K; the potassium absorption of bract and rachis increased by 0.67~1.28 g/plant, 0.35~0.62 g/plant and 0.18~0.30 g/plant (Figure 5).

**Fig 5.**
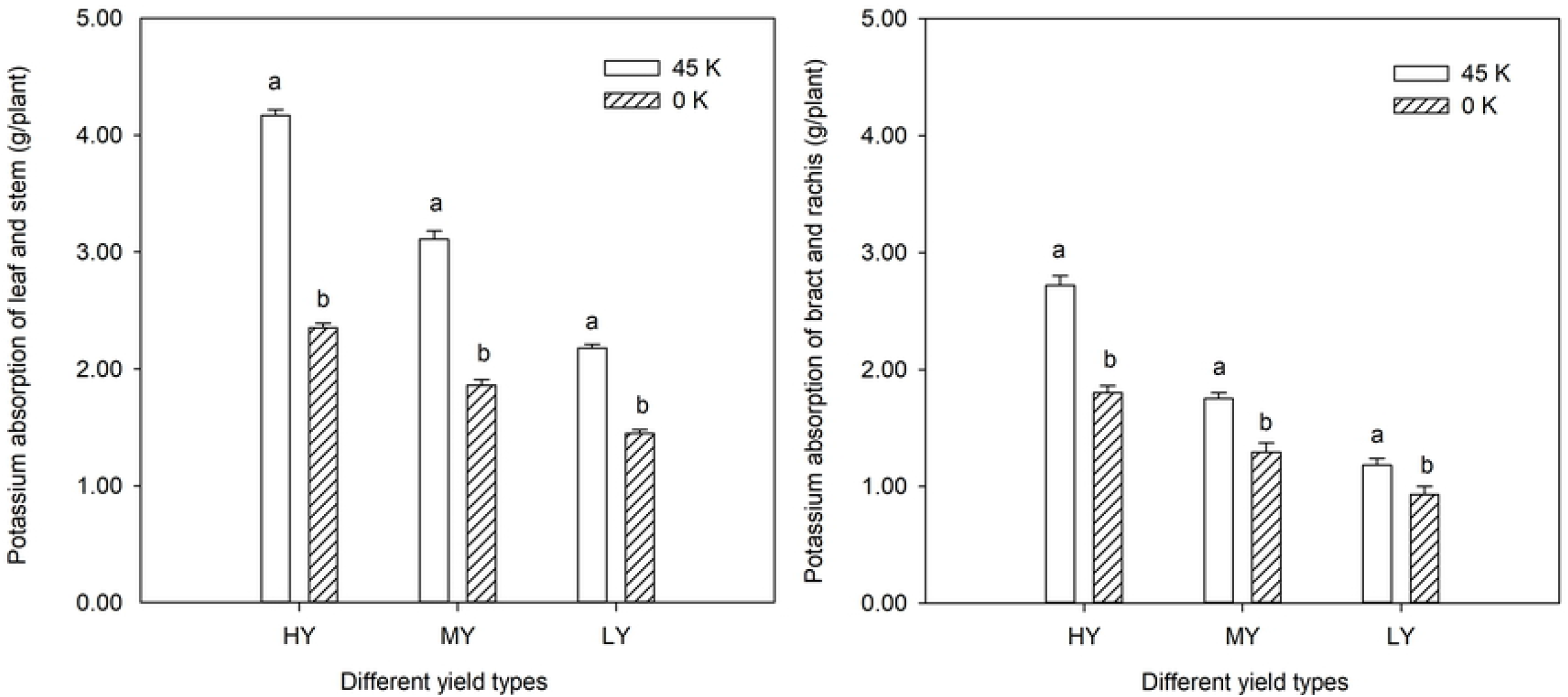
Effects of potassium fertilizer on potassium absorption of maize inbred lines with different yield types in the silking stage

At maturity, the potassium absorption of leaf and stem of high-yield, middle-yield and low-yield type maize inbred lines increased by 2.75~5.44 g/plant, 1.55~1.89 g/plant and 0.62~1.21 g/plant at 45K compared with that at 0K; the potassium absorption of bract and rachis increased by 1.20~2.22 g/plant, 0.74~1.14 g/plant and 0.44~0.65 g/plant; the potassium absorption of grain increased by 1.76~2.65g/plant, 1.17~1.69 g/plant and 0.73~1.13 g/plant (Figure 6).

**Fig 6.**
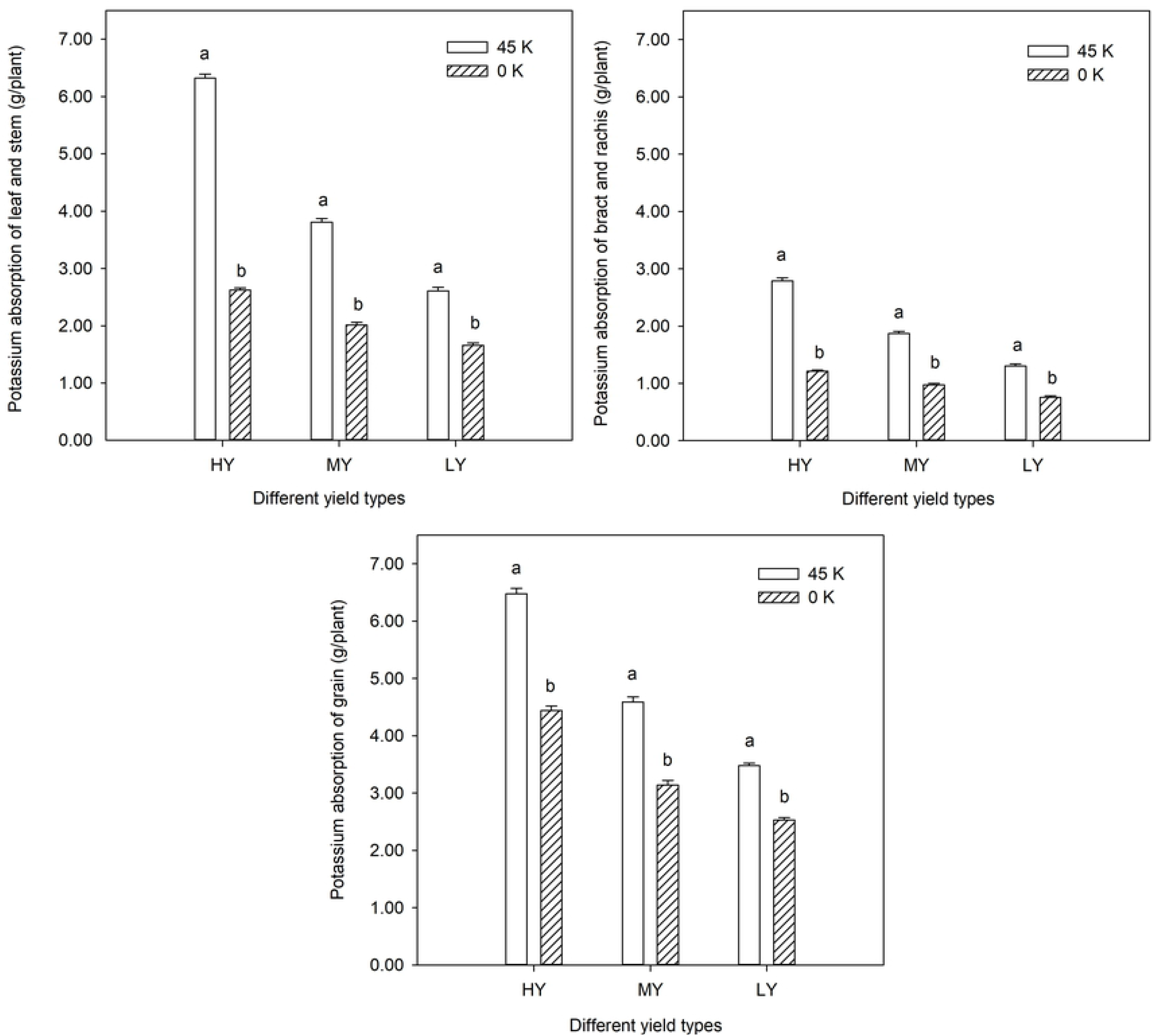
Effects of potassium fertilizer on potassium absorption of maize inbred lines with different yield types at maturity stage

The potassium absorption of maize inbred lines with different yield types were shown as follows: high-yield type>middle-yield type>low-yield type, and potassium fertilizer could significantly increase the potassium absorption of maize.

### Effects of potassium fertilizer on potassium absorption efficiency of maize inbred lines with different yield types

The potassium absorption efficiency of leaf and stem, bract and rachis, grain of maize inbred lines in the silking stage and maturity stage varied significantly (p<0.05) between potassium levels and inbred lines, and the effects of interaction between inbred lines and potassium levels varied significantly (Table 5).

**Table 5.**
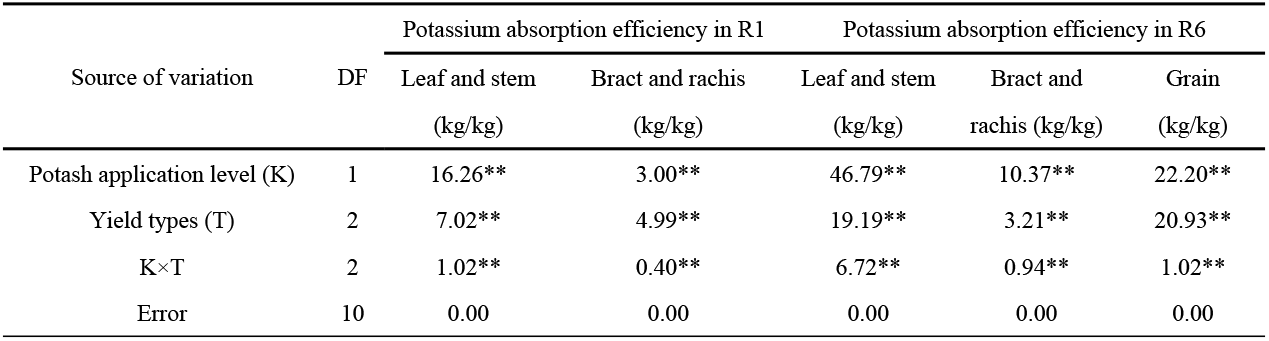
ANOVA results for potassium absorption efficiency of maize inbred lines

| Source of variation | DF | Potassium absorption efficiency in R1 |  | Potassium absorption efficiency in R6 |  |  |
| --- | --- | --- | --- | --- | --- | --- |
|  |  | Leaf and stem<br>(kg/kg) | Bract and rachis<br>(kg/kg) | Leaf and stem<br>(kg/kg) | Bract and rachis<br>(kg/kg) | Grain<br>(kg/kg) |
| Potash application level (K) | 1 | 16.26** | 3.00** | 46.79** | 10.37** | 22.20** |
| Yield types (T) | 2 | 7.02** | 4.99** | 19.19** | 3.21** | 20.93** |
| K×T | 2 | 1.02** | 0.40** | 6.72** | 0.94** | 1.02** |
| Error | 10 | 0.00 | 0.00 | 0.00 | 0.00 | 0.00 |

**Table 6.** Correlation analysis between yield, dry matter and potassium absorption characteristics of maize inbred lines

| Period | Index | R1 |  |  |  |  |  | R6 |  |  |  |  |  |  |  |  |  |
| --- | --- | --- | --- | --- | --- | --- | --- | --- | --- | --- | --- | --- | --- | --- | --- | --- | --- |
|  |  | DMLS | DMBR | PALS | PABR | PAELS | PAEBR | DMLS | DMBR | DMG | PALS | PABR | PAG | PAELS | PAEBR | PAEG | Yield |
|  |  | (g/plant) | (g/plant) | (g/plant) | (g/plant) | (kg/kg) | (kg/kg) | (g/plant) | (g/plant) | (g/plant) | (g/plant) | (g/plant) | (g/plant) | (kg/kg) | (kg/kg) | (kg/kg) | (kg/hm<sup>2</sup>) |
| R1 | DMLS | 1 |  |  |  |  |  |  |  |  |  |  |  |  |  |  |  |
|  | DMBR | 0.981** | 1 |  |  |  |  |  |  |  |  |  |  |  |  |  |  |
|  | PALS | 0.920** | 0.842* | 1 |  |  |  |  |  |  |  |  |  |  |  |  |  |
|  | PABR | 0.985** | 0.963** | 0.949** | 1 |  |  |  |  |  |  |  |  |  |  |  |  |
|  | PAELS | 0.920** | 0.841* | 1.000** | 0.949** | 1 |  |  |  |  |  |  |  |  |  |  |  |
|  | PAEBR | 0.985** | 0.964** | 0.949** | 1.000** | 0.949** | 1 |  |  |  |  |  |  |  |  |  |  |
| R6 | DMLS | 0.974** | 0.993** | 0.862* | 0.974** | 0.861* | 0.974** | 1 |  |  |  |  |  |  |  |  |  |
|  | DMBR | 0.995** | 0.982** | 0.928** | 0.995** | 0.928** | 0.996** | 0.985** | 1 |  |  |  |  |  |  |  |  |
|  | DMG | 0.995** | 0.991** | 0.882* | 0.975** | 0.882* | 0.975** | 0.979** | 0.990** | 1 |  |  |  |  |  |  |  |
|  | PALS | 0.885* | 0.814* | 0.986** | 0.940** | 0.986** | 0.940** | 0.846* | 0.907* | 0.848* | 1 |  |  |  |  |  |  |
|  | PABR | 0.877* | 0.792 | 0.993** | 0.924** | 0.993** | 0.924** | 0.821* | 0.893* | 0.834* | 0.995** | 1 |  |  |  |  |  |
|  | PAG | 0.970** | 0.931** | 0.975** | 0.989** | 0.975** | 0.989** | 0.950** | 0.981** | 0.947** | 0.959** | 0.954** | 1 |  |  |  |  |
|  | PAELS | 0.885* | 0.815* | 0.986** | 0.940** | 0.986** | 0.940** | 0.847* | 0.907* | 0.849* | 1.000** | 0.995** | 0.959** | 1 |  |  |  |
|  | PAEBR | 0.876* | 0.790 | 0.993** | 0.923** | 0.993** | 0.923** | 0.820* | 0.892* | 0.833* | 0.995** | 1.000** | 0.953** | 0.995** | 1 |  |  |
|  | PAEG | 0.970** | 0.931** | 0.975** | 0.989** | 0.975** | 0.989** | 0.950** | 0.981** | 0.946** | 0.959** | 0.954** | 1.000** | 0.959** | 0.953** | 1 |  |
|  | Yield | 0.949** | 0.989** | 0.757 | 0.915* | 0.755 | 0.915* | 0.970** | 0.944** | 0.973** | 0.722 | 0.696 | 0.866* | 0.723 | 0.694 | 0.866* | 1 |

Note: DMLS, DMBR, DMG, PALS, PABR, PAG, PAELS, PAEBR, PAEG indicated the dry matter of leaf and stem, dry matter of bract and rachis, dry matter of grain, potassium absorption of leaf and stem, potassium absorption of bract and rachis, potassium absorption of grain, potassium absorption efficiency of leaf and stem, potassium absorption efficiency of bract and rachis, potassium absorption efficiency of grain respectively.

In the silking stage, the potassium absorption efficiency of leaf and stem of high-yield, middle-yield and low-yield type maize inbred lines increased by 2.23~3.48kg/kg, 1.78~2.15kg/kg and 0.78~1.60kg/kg at 45K compared with that at 0K; the potassium absorption efficiency of bract and rachis increased by 1.00~1.92kg/kg, 0.53~0.92kg/kg and 0.27~0.46kg/kg. The potassium absorption efficiency of leaf and stem of high-yield type maize inbred lines was 1.34 and 1.91 times higher than that of middle-yield and low-yield type at 45K, 1.26 and 1.61 times higher than that of middle-yield and low-yield type at 0K. The potassium absorption efficiency of bract and rachis of high-yield type maize inbred lines was 1.55 and 2.31 times higher than that of middle-yield and low-yield type at 45K, 1.40 and 1.93 times higher than that of middle-yield and low-yield type at 0K (Figure 7).

**Fig 7.**
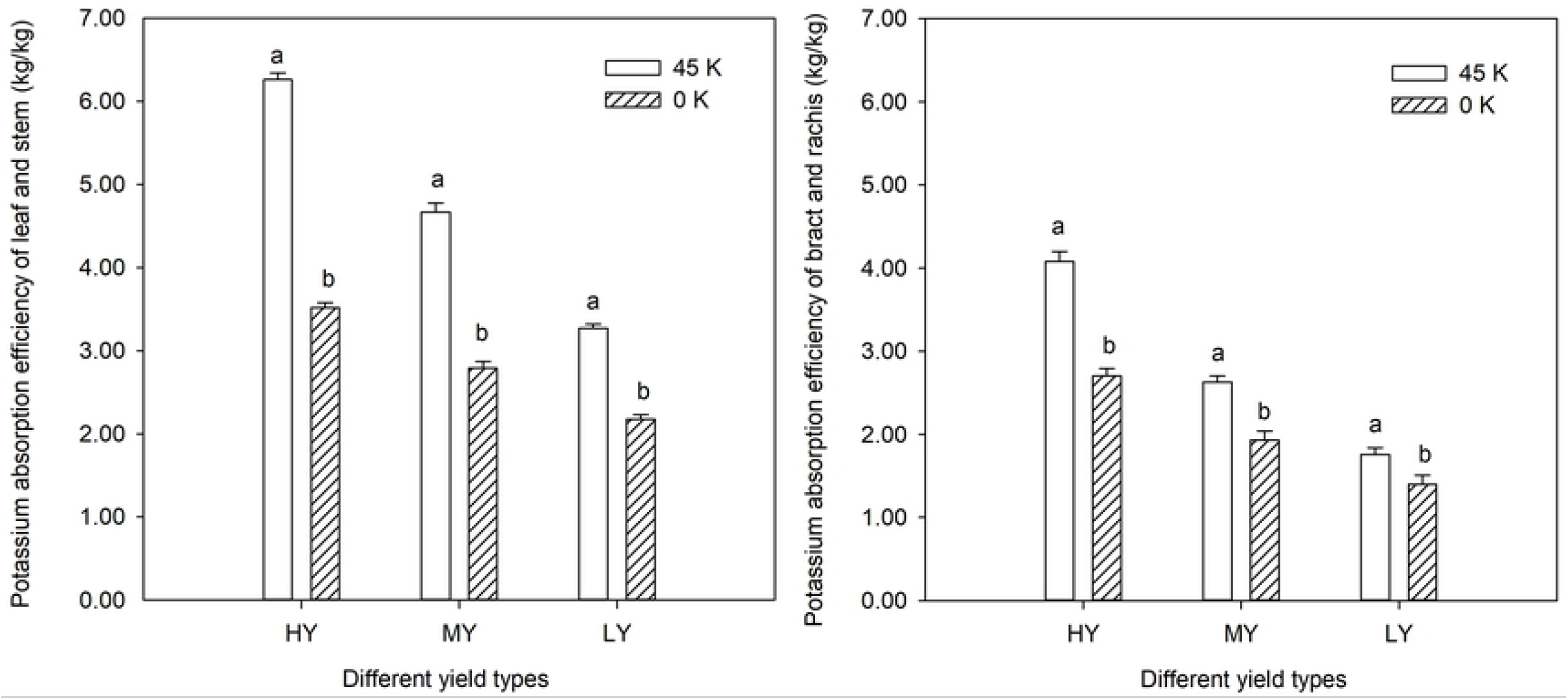
Effects of potassium fertilizer on potassium absorption efficiency of maize inbred lines with different yield types in the silking stage

At maturity, the potassium absorption efficiency of leaf and stem of high-yield, middle-yield and low-yield type maize inbred lines increased by 4.12~8.16kg/kg, 2.33~2.84kg/kg and 0.94~1.81kg/kg at 45K compared with that at 0K; the potassium absorption efficiency of bract and rachis increased by 1.80~3.33kg/kg, 1.10~1.71kg/kg and 0.66~0.97kg/kg; the potassium absorption efficiency of grain increased by 2.64~3.98kg/kg, 1.76~2.53kg/kg and 1.10~1.69kg/kg. The potassium absorption efficiency of leaf and stem of high-yield type maize inbred lines was 1.66 and 2.43 times higher than that of middle-yield and low-yield type at 45K, 1.31 and 1.58 times higher than that of middle-yield and low-yield type at 0K. The potassium absorption efficiency of bract and rachis of high-yield type maize inbred lines was 1.49 and 2.14 times higher than that of middle-yield and low-yield type at 45K, 1.24 and 1.60 times higher than that of middle-yield and low-yield type at 0K. The potassium absorption efficiency of grain of high-yield type maize inbred lines was 1.41 and 1.86 times higher than that of middle-yield and low-yield type at 45K, 1.41 and 1.75 times higher than that of middle-yield and low-yield type at 0K (Figure 8).

**Fig 8.**
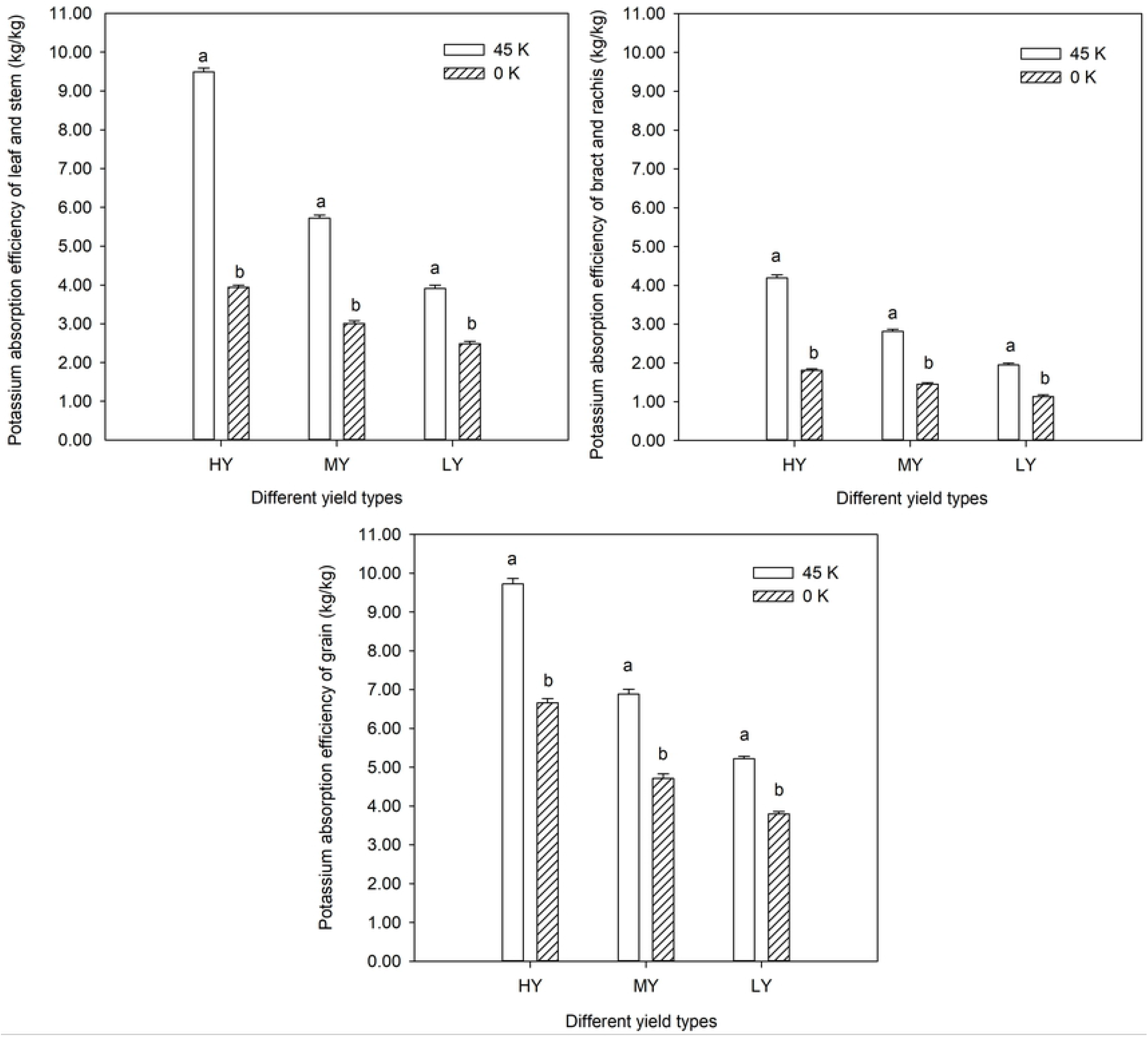
Effects of potassium fertilizer on potassium absorption efficiency of maize inbred lines with different yield types at maturity stage

The potassium absorption efficiency of maize inbred lines with different yield types were shown as follows: high-yield type>middle-yield type>low-yield type, and potassium fertilizer could significantly increase the potassium absorption efficiency of maize.

### Correlation analysis of yield and dry matter, potassium absorption characteristics of maize inbred lines

The dry matter of leaf, stem, bract, rachis of silking stage and yield, the potassium absorption and the potassium absorption efficiency of bract and rachis of silking stage and yield varied positive correlations significantly, and the correlation coefficients were 0.949, 0.989, 0.915 and 0.915, respectively. The dry matter of leaf, stem, bract, rachis, grain of maturity and yield, the potassium absorption and the potassium absorption efficiency of grain of maturity and yield varied positive correlations significantly, and the correlation coefficients were 0.970, 0.944, 0.973, 0.866 and 0.866, respectively.

The results showed that the increase of some plant organs in dry matter, potassium absorption and potassium absorption efficiency could significantly improve the maize yield, including the dry matter of leaf, stem, bract, rachis of silking stage, the potassium absorption and the potassium absorption efficiency of bract and rachis of silking stage, the dry matter of leaf, stem, bract, rachis, grain of maturity, the potassium absorption and the potassium absorption efficiency of grain of maturity.

## Discussion

The cluster analysis method can classify different germplasm resources based on genetic differences among varieties objectively and accurately, so as to screen out excellent germplasm resources and provide basic materials for crop breeding and genetic improvement [19, 20]. It was an important way to achieve stable and high maize yield by researching the yield differences of different genotypes of maize inbred lines and improving maize varieties so as to improve maize yield [21–23]. Therefore, in this study, 20 maize inbred lines were classified into three types by cluster analysis method: high-yield type, middle-yield type and low-yield type. Potassium fertilizer could increase maize yield, the yield of maize inbred lines with high-yield, middle-yield and low-yield type increased by 7.96%~8.45%, 4.72%~5.35% and 2.41%~2.95% at 45K compared with that at 0K. The yield of maize inbred lines with different yield types was as follows: high-yield type>middle-yield type>low-yield type. The test provided basic materials for further research on dry matter accumulation and potassium absorption characteristics of maize inbred lines with different yield types.

The accumulation of dry matter is an important index to improve maize yield [24]. Potassium fertilizer played an important role in promoting the dry matter accumulation of maize [25]. Potassium fertilizer could increase the dry matter of maize leaf, stem, bract, rachis and grain, thus promoting the increase of maize grain yield [26–28]. In this study, the dry matter of maize inbred lines with different yield types increased significantly under potassium application level, and the increase range of high-yield type maize inbred lines was greater than that of middle-yield and low-yield type maize inbred lines, indicating that maize inbred lines with high yield had higher dry matter accumulation ability. Under the condition of 0K, the decrease of dry matter of high-yield type maize inbred lines was smaller. The results showed that high-yield type maize inbred lines had stronger ability to maintain higher dry matter accumulation under low potassium stress. The grain dry matter of maize inbred lines with high-yield, middle-yield and low-yield type increased by 14.29%~15.23%, 9.58%~11.71% and 5.81%~7.63% at 45K compared with that at 0K.

Nutrient absorption is the basis of yield formation, which directly affects the yield and fertilizer absorption and utilization efficiency. The potassium absorption capacity of maize is an important factor to obtain high yield [29]. The absorption capacity of maize plants to potassium could directly affect the accumulation and distribution of dry matter, and then affect the formation of maize yield [30, 31]. In this study, there were significant differences in potassium absorption and potassium absorption efficiency of maize inbred lines in the silking stage and maturity stage under the two potassium fertilizer levels, which were as follows: high-yield type>middle-yield type>low-yield type. Potassium application increased the amount and efficiency of potassium absorption. The improvement degree of potassium absorption characteristics of high-yield type maize inbred lines was greater than that of middle-yield and low-yield type maize inbred lines. The grain potassium absorption of maize inbred lines with high-yield, middle-yield and low-yield type maize inbred lines increased by 1.76~2.65g/plant, 1.17~1.69 g/plant and 0.73~1.13 g/plant at 45K compared with that at 0K, the potassium absorption efficiency increased by 2.64~3.98 kg/kg, 1.76~2.53 kg/kg and 1.10~1.69 kg/kg. The results showed that high-yield type maize inbred lines were more sensitive to potassium fertilizer in improving potassium absorption characteristics, which may be one of the important factors for high-yield type maize inbred lines to gain higher yield. The correlated relationship between maize yield and dry matter accumulation and potassium absorption characteristics was of great significance to guide the breeding of maize materials with high yield and good quality.

## Conclusions

20 maize inbred lines were divided into three types, the first type was high-yield type, including FAPW, PHG39, LH51, K12, Shen 5003 and NP49; the second type was middle-yield type, including Huang C, Jun 92-8, Zi 330, P2, LP5, 7810 and IRF233; the third type was low-yield type, including IB014,Ji 63, G303, IRF236, M3401, L269 and DH382. It provided the basic material for the future research on the physiological characteristics of maize inbred lines of different yield types, and provided the suitable maize inbred materials for the rational utilization of potash resources.

Potassium fertilizer can improve the dry matter accumulation capacity and potassium absorption efficiency, influence the growth and development of maize, and then improve the grain yield. High-yield type maize inbred lines have stronger ability to maintain higher dry matter accumulation and potassium absorption characteristics under low potassium stress, which can be one of the important factors for high yield of high-yield type maize inbred lines.

## Acknowledgments

We would like to thank the Maize High-Yield and High-Efficiency Cultivation Team for field and data collection.

## Author Contributions

Funding acquisition: Julin Gao, Jiying Sun.

Investigation: Yafang Fan.

Methodology: Jiying Sun, Jian liu, Zhijun Su, Shuping Hu, Zhigang Wang, Xiaofang Yu.

Conceptualization: Julin Gao, Jiying Sun, Jian liu, Yafang Fan.

Writing – original draft: Yafang Fan, Jian liu.

Writing – review & editing: Julin Gao, Jiying Sun.

